# Resistance to diet induced obesity in the apolipoprotein E deficient mouse is associated with an attenuated transcriptional response in visceral fat

**DOI:** 10.1101/494195

**Authors:** Timothy P. Fitzgibbons, Mark Kelly, Jason K. Kim, Michael P. Czech

## Abstract

The apolipoprotein E knockout (EKO) mouse is a well-established model of atherosclerosis. Macrophages in the arterial intima of EKO mice serve a protective role, scavenging oxidatively modified LDL in order to protect cells from toxic free cholesterol. Recent studies have highlighted a similar role for macrophage foam cells in restraining the increased rates of lipolysis in adipose tissue of obese and fasting mice. Interestingly, EKO adipocytes have been shown to have increased rates of lipolysis *in vitro*. Therefore, the aim of this study was to examine how apoE deficiency might alter the transcriptional response of visceral adipose tissue (VAT) to high fat diet (HFD). EKO mice fed HFD for 24 weeks gained less fat mass and were more insulin sensitive than their wild type (WT) littermates. Metabolic cages showed that HFD EKO mice had increased post-prandial oxygen consumption and increased serum β-hydroxybutyrate. DNA microarrays revealed that EKO VAT was comparatively insensitive to HFD in terms of alterations in gene expression, with only 0.1% of probe sets differentially expressed. In contrast, the VAT of WT mice had a 30 fold more extensive alteration in gene expression (3% of probes sets), characterized predominantly by increased expression of immune cell specific genes. In addition, analysis of *a priori* determined gene sets revealed broad down-regulation of PPARγ target and fatty acid catabolism genes in WT VAT, and increased expression of lipid storage and cholesterol synthesis genes. In comparison, expression of PPARγ target genes was not down-regulated in EKO VAT and expression of fatty acid oxidation genes was increased. In summary, we report three novel findings with regards to metabolism in the EKO mouse: 1) increased post-prandial oxygen consumption, 2) increased serum β hydroxybutyrate concentrations and 3) a dramatically less robust transcriptional response to HFD in EKO VAT. These findings suggest that limiting adipocyte exposure to dietary fatty acids may be an attractive therapy for diet induced obesity, provided that compensatory mechanisms that prevent hyperlipidemia can be activated.

## Introduction

Apolipoprotein E (apoE) is a 34 kD glycoprotein which is a component of all lipoprotein particles with the exception of low density lipoprotein (LDL)^1^. It is a ligand for the clearance of chylomicrons and intermediate density lipoprotein particles (IDL) from the circulation via the low density lipoprotein receptor (LDLR), very low density lipoprotein receptor (VLDLR) and low density lipoprotein related protein 1(LRP1)^2^. Apolipoprotein E knockout (EKO) mice have dramatic elevations of plasma cholesterol (5 fold normal) and reduced high density lipoprotein (HDL) concentrations ^3, 4^. These findings mimic those of human patients with Type III hyperlipoproteinemia, which results in increased plasma triglyceride (TG) and cholesterol and premature atherosclerosis.

Apoe is highly expressed in the liver, adipose tissue, macrophages, and other tissues of humans and mice. In adipocytes, apoE expression is increased by PPARγ agonists and decreased by TNF-α^5^. Multiple recent studies have documented protection of the EKO mouse from diet induced obesity (DIO) and insulin resistance^6–9^. Whole body deficiency of apoE also prevents weight gain and insulin resistance in genetic models of obesity such as the *ob/ob* or *Ay*/+ mouse^7, 9, 10^. Protection from diet induced obesity can be reversed by adenoviral over expression of apoE in the liver, suggesting an important role for circulating apoE in adipose tissue lipid uptake^7^.

On the other hand, endogenous adipocyte apoE expression is required for the uptake and storage of free fatty acids. For example, primary adipocytes from EKO mice cultured *in vitro* synthesize less triglyceride (TG) than WT when incubated with VLDL^11^. Furthermore, EKO adipocytes transplanted into WT mice fail to increase in size in response to 10 weeks of HFD^11^. There are at least two mechanisms by which absence of apoE impairs TG synthesis in adipocytes^2, 11^. Binding and whole particle uptake of VLDL to EKO adipocytes is impaired due to reduced expression and/or decreased cell surface localization of VLDLR, LDLR, and LRP1^11^. Lipoprotein lipase (LPL) dependent TG synthesis is also decreased in EKO adipocytes. This is thought to be due impaired fatty acid translocation across lipid rafts, as caveolin 1 expression is markedly reduced in EKO adipocytes^11^. These same mechanisms may also account for reduced lipid uptake in liver and skeletal muscle, protection from lipotoxicity, and preservation of whole body insulin sensitivity despite severe hyperlipidemia^2, 12^.

Macrophage uptake of triglyceride rich lipoproteins (TRLP), unlike adipocytes, is not dependent on apoE. Foam cells take up modified lipoproteins via scavenger receptors, predominantly SRA and CD36^13^. Rather, apoE in macrophages is important in reverse cholesterol transport and lipidation of ApoA1 and HDL by cholesterol export via ABCA1 and ABCG1^14^. Two studies have shown that bone marrow transplantation of apoE^+/+^ macrophages into EKO mice ameliorates atherosclerosis; this is accompanied by improvements in serum lipoprotein profiles to wild type levels^15, 16^. Therefore, in contrast to adipocytes and other tissues, macrophage specific apoE appears to serve a protective effect in respect to lipid handling^14^. This serves to distinguish the role of the macrophage in atherosclerosis from its role in diet induced obesity, the depletion of which, improves metabolic disease. The specific lipid moieties that stimulate foam cell formation in each case might account for these differences. While it is well established that modified lipoprotein particles are the stimulus for foam cell formation in atherosclerosis, it has only recently been suggested that free fatty acids may be responsible for foam cell formation in circumstances of increased adipocyte lipolysis, such as obesity or fasting^17^. Interestingly, adipocytes from EKO mice have been shown to have increased rates of lipolysis *in vitro*, thus we would expect greater inflammation in the VAT from these mice^5^.

Therefore, the purpose of this study was to determine 1) how does adipocyte gene expression change in response to HFD the absence of endogenous adipocyte apoE? and 2) is the visceral adipose tissue of EKO mice characterized by increased foam cell accumulation analogous to the atherosclerotic plaque?

## Materials and Methods

### Animal Studies

An original colony *Apoe*^*-/-*^ (*Apoe^tm1Un^*) on the C57BL6/J background was purchased from the Jackson Laboratory (Bar Harbor, ME). Mice were backcrossed to C57BL6/J to obtain WT control littermates. All animals were fed normal chow until 8 weeks of age. Male wild type and EKO mice were then divided into two groups (n=5-6 per group); one fed normal chow and one fed Western diet (42% Milk fat, 0.2% cholesterol, Harlan TD88137) for 24 or 38 weeks respectively. Animals were fed *ad libitum* with free access to water and housed in the University of Massachusetts (UMASS) Medical School Animal Facility with a 12:12 hour light-dark cycle. Animals were weighed weekly for the duration of the diet study. Intraperitoneal glucose tolerance testing was performed as previously described ^18^. Area under the curve for the GTT was calculated using the trapezoidal method. At the completion of HFD, mice were fasted for 4 hours and then euthanized with CO2 inhalation and bilateral pneumothorax. Livers and epididymal VAT were then harvested and snap frozen in liquid nitrogen. All experiments were performed in accordance with protocols approved by the Animal Care and Use Committee at UMASS Medical School.

### Metabolic Phenotyping

Body composition was determined non-invasively in awake mice using ^1^H-MRS (Echo Medical System). For metabolic studies, mice were housed under controlled temperature and lighting with free access to food and water. The food/water intake, energy expenditure, respiratory exchange ratio, and physical activity were performed on three consecutive days using metabolic cages (TSE Systems). Post prandial oxygen consumption was calculated by subtracting the pre-fed VO2 (ml/hr)(16:00) from the post-fed (VO2)(6:00) as previously described^19^.

### Free Fatty Acid and β hydroxybutyrate Levels

Blood was collected via cardiac puncture at the time of euthanasia into heparin coated micro centrifuge tubes and centrifuged at 3000 x g for 15 minutes. Serum was then decanted and stored at -80 °C. Serum free fatty acids were measured with the NEFA (2) colorimetric kit according the manufacturer’s instructions (Wako Diagnostics, Richmond, VA). β hydroxybutyrate was measured with a colorimetric assay (#700190) according to the manufacturer’s instructions (Cayman Chemicals, Ann Arbor, MI).

### Quantitative PCR

Adipose tissue was isolated as previously described, snap frozen in liquid nitrogen and stored at -80°C. Tissues were homogenized and total RNA was isolated with RNA Mini Lipid kits (Qiagen, Valencia, CA). 250 ng total RNA was reverse transcribed with the Bio-Rad iScript cDNA Synthesis Kit (Bio-Rad, Hercules, CA). cDNA was diluted 1:5 and 2.5 μl was used in a 12.5 μl reaction volume; each reaction was performed in duplicate. Real time qPCR was performed with a Bio-Rad C1000 Thermal Cycler and SYBR Green Master Mix (Bio-Rad, Hercules, CA) using the following cycle parameters: 95.0° C for 3 min, 95.0° for 0:10 min, 60.0° C for 0:15 min, 72.0° C for 0:30 min for 40 cycles. Expression was normalized to the reference gene *36b4* and expressed as relative to expression in VAT of normal diet mice of the same genotype using the 2 ^−ΔΔct^ method ^20^. Melt curve analysis was performed to determine the specificity of the PCR reaction products.

### Microarray Analysis

RNA was isolated from epididymal VAT as previously described. RNA concentrations were determined using a Nanodrop 2000 Spectrophotometer (Thermo Fisher, Willmington, DE). The RNA quality was assessed using an Agilent 2100 Bioanalyzer (Agilent Technologies, Santa Clara, CA). Only samples with a RNA Integrity Number >7.5 and normal 18 and 28s fractions on microfluidic electrophoresis were used. 250 ng total RNA was used as template for cDNA synthesis and *in vitro* transcription using the Ambion WT Expression kit (Ambion, Carlsbad, CA). Second strand cDNA was then labeled with the Affymetrix WT Terminal Labeling kit and samples were hybridized to Affymetrix Mouse Gene 1.0 ST arrays (Affymetrix, Santa Clara, CA). Gene chip expression array analysis for individual genes was performed as previously described ^18^, filtering for p <0.05 and a fold change of >1.5. Four biological replicate hybridizations per tissue and diet were performed, for a total of 16 hybridizations. Robust multi-array average (RMA) was adopted in the UMASS Microarray Computational Environment (MACE) to preprocess raw oligonucleotide microarray data. The algorithm was implemented as a function of the R package Affy ^21^, which is part of the Bioconductor project ^22^ using the statistical computing language R (R Foundation for Statistical Computing, Vienna, Austria). All statistical calculations are performed using the R statistical computing environment and results are stored in a relational database. The preprocessed data are stored as base 2 log transformed real signal numbers and are used for fold-change calculations and statistical tests and to determine summary statistics. Mean signal values and standard deviations are computed for each gene across triplicate experiments and stored in the database. The fold change of expression of a gene in two experiments is the ratio of mean signal values from these experiments and is always a number greater than one. If the ratio is less than one, the negative value of the inverse ratio is stored as fold change. All down regulated genes therefore have a negative fold change value, up regulated genes have a positive fold change. In both cases this value is greater or equal than one.

To determine differential expression of genes in two hybridization experiments MACE internally conducts a Student’s t-test with the expression signal values of the two hybridizations for all genes in the set. The t-test value and test p-value are stored in MACE and can be queried through the MACE user interface. The p-values stored and displayed as a result of a query are not adjusted for multiple testing.

### Histology and Immunohistochemistry

Adipose tissue samples (n=3 per group) from normal and high fat diet animals were fixed in 4% formalin for immunohistochemistry. Briefly, samples were embedded in paraffin, sectioned, and stained with a rat anti-mouse F4/80 (ABd Serotec, Raleigh, NC)(1:40 dilution). Staining was visualized with a HRP linked rabbit anti-rat secondary anti-body. Staining with the secondary antibody alone was performed as a negative control. Images were taken with a Zeiss Microscope and PixeLINK SE Software.

### Statistical Analysis

All values are shown as mean ± SEM. For experiments other than the microarray analyses, the Student’s *t* test for two tailed distributions with equal variances was used for comparison between 2 groups; for comparisons greater of ≥ 3 groups, two way ANOVA followed by the Bonferroni correction was used. Differences less than p < 0.05 were considered significant. Data was entered into Microsoft Excel and statistical analyses were performed with Graph Pad Prism 5.

## Results

### EKO mice are resistant to DIO after 24 and 38 weeks of HFD

Male WT and EKO mice were started on normal diet (ND) or HFD at 8 weeks of age (n=5-6 per group). After 24 weeks, wild type HFD mice were significantly more obese than their ND littermates (48.0 ± 1.6 vs. 33.0 ± 0.8 gms, p<0.001)(Figure 1A). In contrast, there was no difference between ND and HFD fed EKO mice (34.9 ± 0.5 vs. 34.8 ± 1.1 gms, p=NS). Body composition analyses of mice at 24 weeks of HFD revealed a fivefold increase in fat mass of WT mice, whereas HFD EKO mice had a non significant doubling of fat mass (Table 1). Diet and genotype had no influence on lean body mass (Table 1).

**Figure 1.**
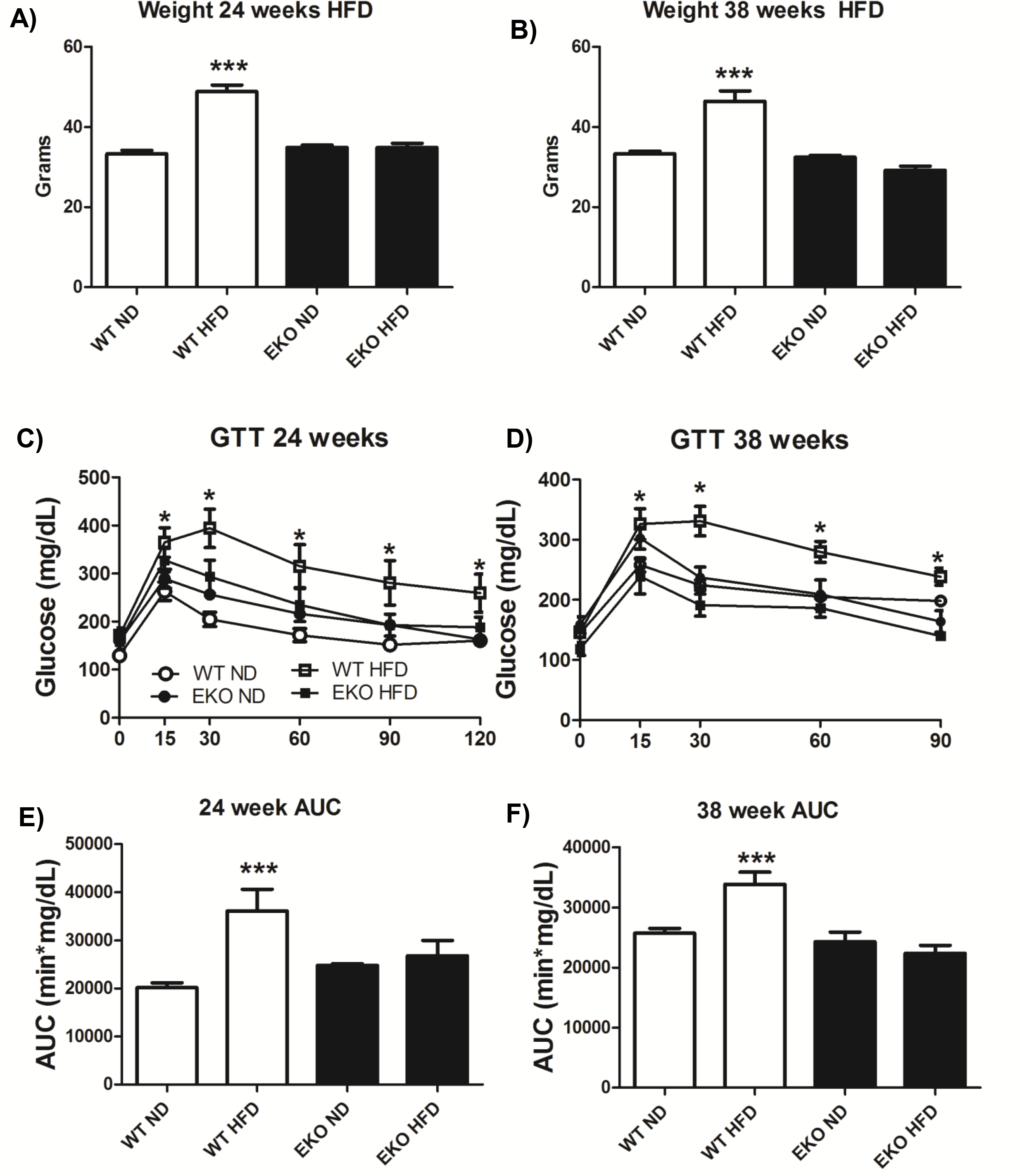
EKO mice are resistant to diet induced obesity and impaired glucose tolerance after 24 and 38 weeks of HFD. A,B) Weight of WT and EKO mice after 24 and 38 weeks of HFD (n=5-6 per group). C,D) Intraperitoneal GTT in WT and EKO mice after 24 and 38 weeks of HFD. E,F) Area under the curve for the IP GTT at 24 and 38 weeks. *p< 0.05 vs. WT ND, ***p<0.001 vs. WT ND using the Student’s t-test

**Table 1.** Metabolic Parameters of WT and EKO Mice fed ND and HFD for 24 Weeks.

|  | WT ND (n=5) | WT HFD (n=5) | EKO ND (n=5) | EKO HFD (n=6) |
| --- | --- | --- | --- | --- |
| Weight (gms) | 31.4 ± 0.7 | 41.9 ± 0.3*** | 32.0 ± 1.0 | 35.0 ± 2.3 |
| Fat mass (gms) | 3.6 ± 1.0 | 15.3 ± 0.2*** | 3.1 ± 0.7 | 7.0 ± 1.9 |
| Lean mass (gms) | 26.9 ± 0.8 | 25.8 ± 0.3 | 28.3 ± 0.7 | 27.3 ± 0.8 |
| 24 Hour VO<sub>2</sub> Consumption (ml/hr/kg) | 3512 ± 126 | 3073 ± 24 | 3671 ± 126 | 3804 ± 294* |
| Light phase VO<sub>2</sub> Consumption (ml/hr/kg) | 3424 ± 116 | 2883 ± 57 | 3470 ± 38 | 3670 ± 267** |
| Dark phase VO<sub>2</sub> Consumption (ml/hr/kg) | 3801 ± 142 | 3263 ± 21 | 4001 ± 150 | 4066 ± 333 * |
| 24 Hour VCO<sub>2</sub> Production (ml/hr/kg) | 3038 ± 138 | 2379 ± 49 | 3112 ± 143 | 2988 ± 251* |
| 24 Hour Energy Expenditure Rate (kcal/hr/kg) | 17.6 ± 0.65 | 14.7 ± 0.13 | 17.9 ± 0.65 | 18.3 ± 1.43 |
| Raw 24 Hour VO<sub>2</sub> Consumption (ml/hr) | 112.8 ± 3.2 | 128.0 ± 1.05 | 117.2 ± 3.3 | 125.8 ± 2.8 |
| Respiratory Exchange Ratio (RER) | 0.8 ± 0.01 | 0.7 ± 0.01††† | 0.8 ± 0.01 | 0.7 ± 0.01††† |
| Food intake (gms/day) | 3.6 ± 0.55 | 2.3 ± 0.33 | 4.3 ± 0.47 | 2.7 ± 0.14 |
| Physical activity (counts/day) | 4853 ± 413 | 4853 ± 456 | 5886 ± 1219 | 4717 ± 726 |
| Free fatty acids (mmol/L) | 0.5 ± 0.05 | 0.8 ± 0.16 | 2.2 ± 0.35<sup>^^</sup> | 3.1 ± 3.1<sup>^^</sup> |
| Fasting plasma glucose (mg/dl) | 185 ± 13 | 204 ± 11 | 146 ± 14 | 148 ± 13* |
| GTT AUC (min*mg/dl) | 32367 ± 2724 | 44748 ± 3384<sup>***</sup> | 26532 ± 2976 | 32735 ± 2352 |
| β hydroxybutyrate (mM) | 0.36 ± 0.08 | 0.83 ± 0.24 | 3.0 ± 0.24 | 4.9 ± 0.56<sup>†</sup> |

\*\*\*p<0.001 vs. WT ND using the Student's t-test,
\*p<0.05, \*\*p<0.01 vs. WT HFD using two way ANOVA and the Bonferroni correction
††† p<0.001 vs. ND within genotype using two way ANOVA and the Bonferroni correction
<sup>^^</sup>p<0.001 vs. WT ND, HFD using two way ANOVA and the Bonferroni correction.
† p<0.001 vs. EKO ND using two way ANOVA and the Bonferroni correction.

Fasting plasma glucose was significantly lower in HFD EKO than HFD WT controls and free fatty acids were significantly higher (Table 1). Intraperitoneal GTT revealed that HFD WT mice had impaired glucose tolerance compared to their ND controls at all time points (p<0.05)(Figure 1C). In contrast, there was no significant difference in glucose tolerance between ND and HFD fed EKO mice.

In a second cohort of animals, we decided to extend the duration of HFD to determine if age related impairments in insulin sensitivity might influence fat accumulation in EKO mice. Surprisingly, even after 38 weeks of HFD, there was no difference in weight between ND and HFD fed EKO mice (32.4 ± 0.44 vs. 30.20 ± 0.54, p=NS), nor were there differences in glucose tolerance as seen by IP GTT (Fig 1D, F).

In summary, 24 and 38 weeks of a diet high in saturated fat and cholesterol results in obesity and diabetes in WT but not EKO mice. Increased body mass is due to a five-fold increase in fat mass of HFD fed WT mice. This is consistent with prior studies and extends these data to high fat feeding of chronic duration.

### Metabolic cage studies reveal no differences in energy expenditure, food intake, or physical activity

We hypothesized that EKO mice might have increased energy expenditure or decreased food intake which would account for failure to increase their fat mass with high fat feeding. Metabolic cage analyses were performed for 3 consecutive days after mice had been fed HFD for 23 weeks. There were no differences in food intake or physical activity between genotypes; HFD mice tended to eat less than their normal chow controls (Table 1). Likewise, there were no differences in 24 hour energy expenditure rates (Table 1).

### EKO Mice have increased post-prandial VO2

EKO HFD fed mice had significantly greater rates of normalized VO2 over a 24 hour period than WT HFD fed mice (3804 ± 294 vs. 3073 ± 24 ml/hr/kg, p<0.05)(Table 1)(Figure 2A). Increased rates of VO2 in EKO mice were present in both the light and dark phases. However, when non-normalized VO2 rates were compared, there was no genotype effect, but there was an effect of HFD to increase VO2 (ml/hr)(Figure 2B)(Table 1). ANCOVA confirmed that there was no statistically significant relationship between genotype and VO2 or EE (Table 2). In contrast, HFD increased VO2 (R=0.58, p<0.05) and EE (R=0.76, p<0.001) in both genotypes (Table 2, Supp. Fig. 2).

**Figure 2.**
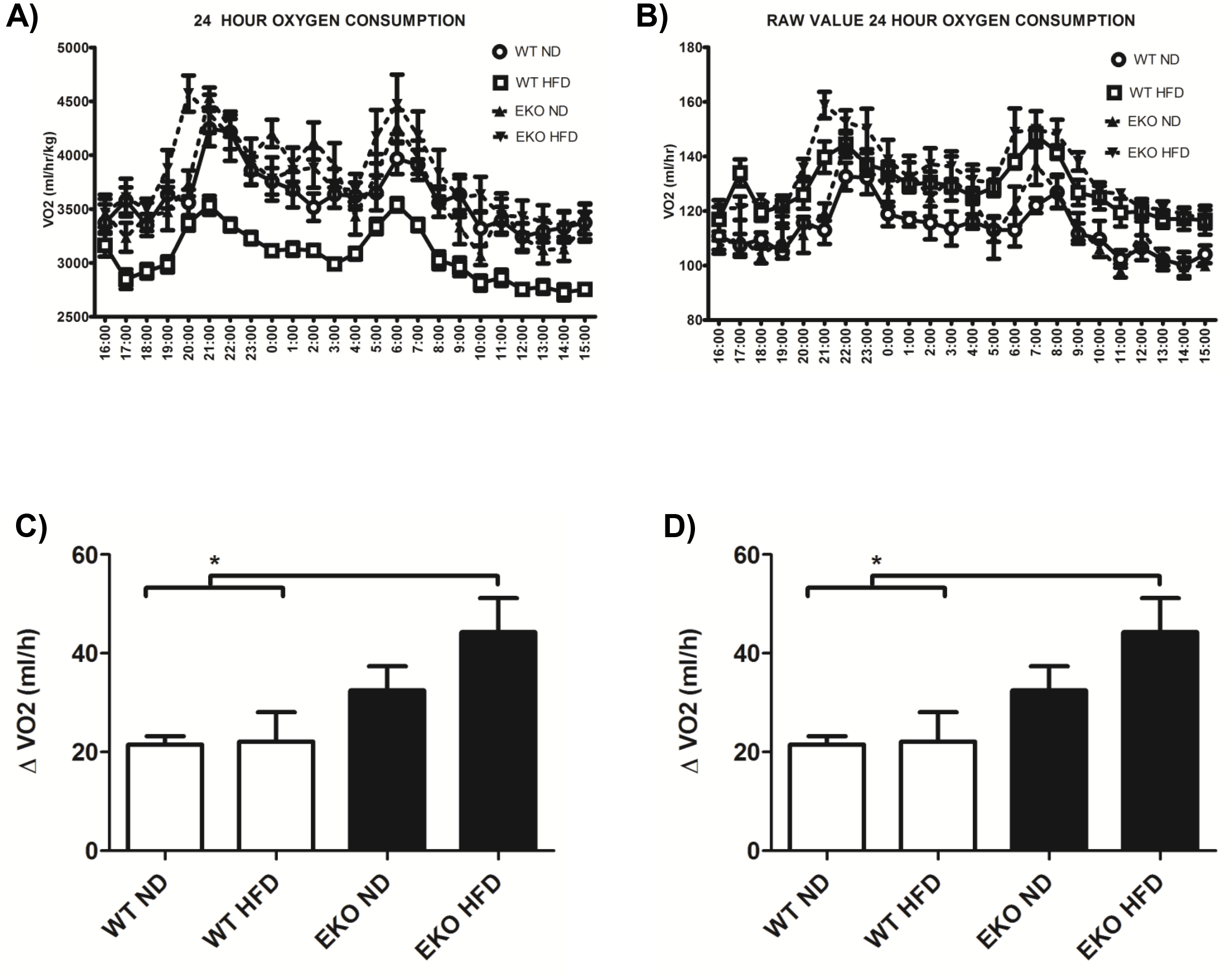
Post prandial, but not 24 hour VO2, is increased in EKO mice. (A) VO2 (ml/hr/kg) was measured using metabolic cages over three consecutive days after 23 weeks of HFD in WT and EKO mice (n=5-6 per group). EKO HFD mice had significantly greater rates of VO2 over 24 hours than WT HFD mice when normalized by weight. (B) When normalization by weight is removed, there is no effect of genotype on total 24 VO2 (ml/hr). (C) The difference in VO2 (ml/hr/kg) before and after the dark (feeding) cycle. The difference VO2 was significantly greater in EKO HFD than WT ND and WT HFD fed mice. (D) The difference in post-prandial VO2 (ml/hr) remained when normalization by weight was removed. *<0.05 vs. WT ND, WT HFD using two way ANOVA and the Bonferroni correction.

**Table 2.** ANCOVA Results for EE and VO2

| Model for EE | Source | DF | F Value | P value |
| --- | --- | --- | --- | --- |
|  | Genotype | 1 | 1.93 | 0.186 |
|  | Diet | 1 | 29.08 | <0.001* |
|  | Genotype * Diet | 1 | 5.32 | 0.036* |
|  | Weight | 1 | 0.11 | 0.746 |
|  | Fat Mass | 1 | 0.01 | 0.942 |
|  | Lean Body Mass | 1 | 0.71 | 0.413 |
| Model for VO2 | Source | DF | F Value | P Value |
|  | Genotype | 1 | 0.36 | 0.560 |
|  | Diet | 1 | 8.41 | 0.011* |
|  | Genotype * Diet | 1 | 1.87 | 0.193 |
|  | Weight | 1 | 0.16 | 0.699 |
|  | Fat Mass | 1 | 0.11 | 0.749 |
|  | Lean Body Mass | 1 | 0.33 | 0.572 |
| *Diet is a significant independent predictor of EE and VO2 using ANCOVA. |  |  |  |  |

As it has been shown that EKO mice have a prolonged clearance time of dietary TG, we hypothesized that they might have an increased post-prandial VO2 consumption. Indeed, the difference between pre-prandial (16:00 hours) and post-prandial (6:00 hours) VO2 (ml/hr/kg) was significantly greater in EKO HFD than WT mice on both diets (p<0.05)(Figure 2C). This effect remained significant when non-normalized VO2 (ml/hr) was used (Figure 2D).

### EKO mice are protected against hepatic steatosis induced by 24 weeks of HFD

Although it is well known that EKO mice have severe elevations of blood cholesterol and TG, given that they have greater rates of VO2 (ml/hr) in the post-prandial period, we hypothesized that they may be consuming some fraction of this excess fatty acid via hepatic β oxidation. Light microscopy showed that HFD EKO mice were protected from hepatic steatosis, while HFD fed WT mice had severe hepatosteatosis with hepatocyte ballooning and large lipid droplet formation (Figure 3B). Indeed, WT HFD mice had a 5 fold increase in liver triglyceride concentrations compared to WT ND mice, whereas EKO HFD had no such increase in liver triglyceride in comparison to their ND controls (Figure 3E). Finally, mRNA expression of the lipid droplet protein genes *Cidea* and *Fsp27* was increased 1000 and 100 fold respectively in WT HFD fed mice (p<0.05)(Figure 3F).

**Figure 3.**
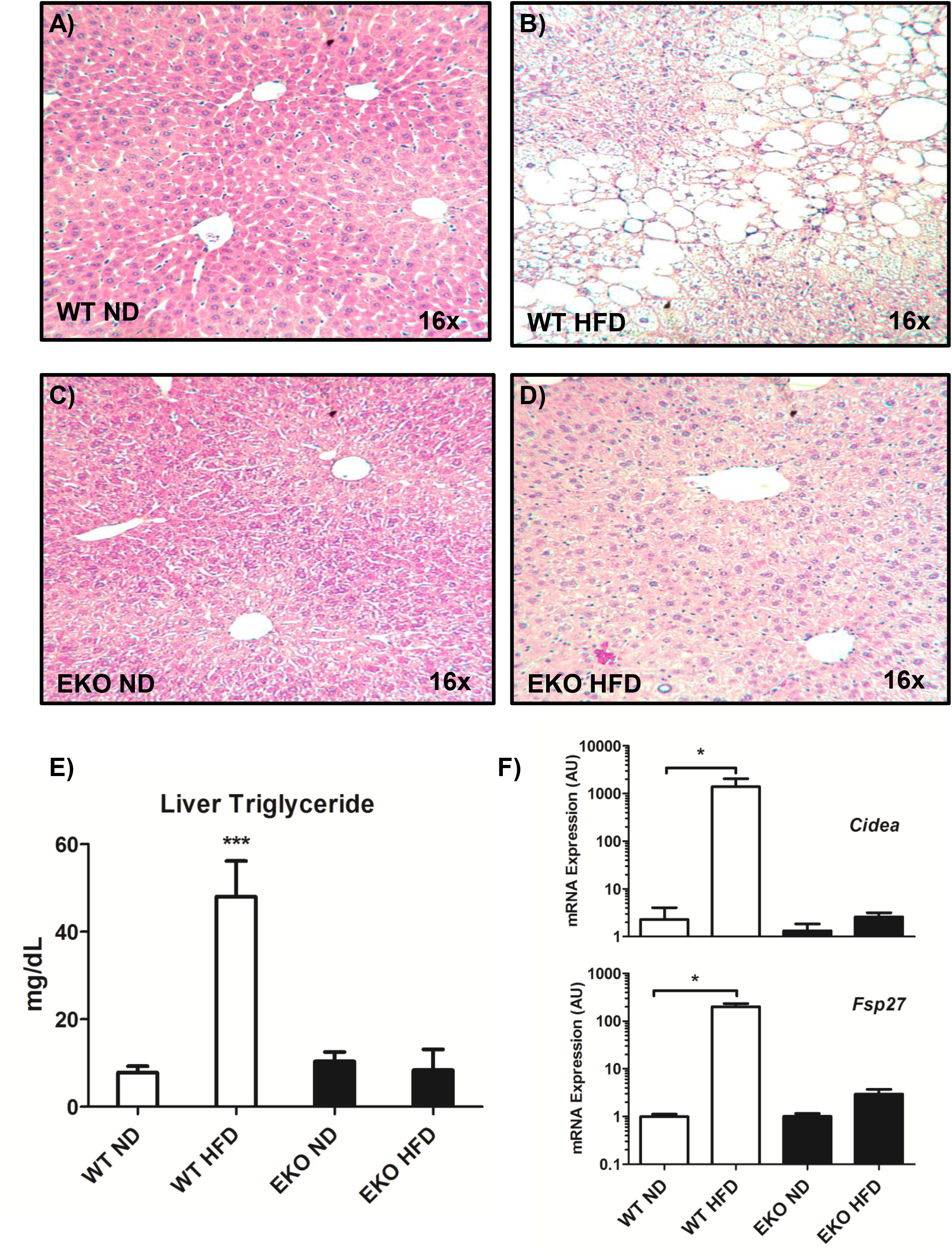
EKO mice are protected against hepatic steatosis induced by 24 weeks of HFD. (A-D) Hematoxylin & eosin staining of liver sections from WT (top) and EKO mice (bottom) after 24 weeks of ND (left) or HFD (right). WT HFD mice have severe hepatocyte ballooning and macrosteatosis. In contrast, EKO HFD mice have much less lipid accumulation and hepatocyte ballooning. (E) Liver triglyceride content in WT ND, WT HFD, EKO ND, and EKO HFD mice (n=6 per group). ***p<0.001 vs. WT ND, EKO ND, and EKO HFD using two way ANOVA and the Bonferroni correction. (F) qPCR expression of the lipid droplet protein genes *Cidea* and *Fsp27* is increased in WT HFD fed mice but not EKO HFD fed mice. *p<0.05 vs. WT ND using the Student’s T test.

Although liver TG concentrations were significantly lower in HFD fed EKO mice, serum free fatty acids were greater in ND and HFD fed EKO mice (Table 1). Serum β hydroxybutyrate concentrations were also 5 fold higher in EKO HFD than WT HFD mice (4.9 ± 0.56 vs. 0.83 ± 0.24 mM, p<0.001)(Table 1). This supported our hypothesis that decreased fat storage in EKO adipose tissue results in increased delivery of fatty acids to the liver, which may be used as substrate for β oxidation and ketone formation.

### Comparative insensitivity of EKO VAT gene expression in response to HFD

Given that EKO mice have an impaired ability to uptake and store dietary fat as triglyceride, we hypothesized that EKO adipocytes might compensate by stimulating *de novo* lipogenesis. Increased glucose uptake by adipose tissue would allow for this and partially explain the improved glucose tolerance of EKO mice. Therefore, we queried expression of key transcription factors (*Srebf1, Nr1h3/Lxrα*) and regulatory enzymes (*Acly, Fasn*) which regulate *de novo* lipogenesis using qPCR. We were surprised to find that obese HFD WT mice had significantly increased expression of *Fasn* (*3.3 fold*)*, Acly* (*4.3 fold*), *Srebf1* (*3.0 fold*), and *Dgat2* (*5.9 fold*) while HFD EKO mice had no such increase (Figure 4). In comparison to ND controls, HFD fed EKO mice had a decrease in expression of all these genes, two of which were statistically significant (Lxrα/*Nr1h3 0.44 fold, Dgat2 0.33 fold*)(Figure 4).

**Figure 4.**
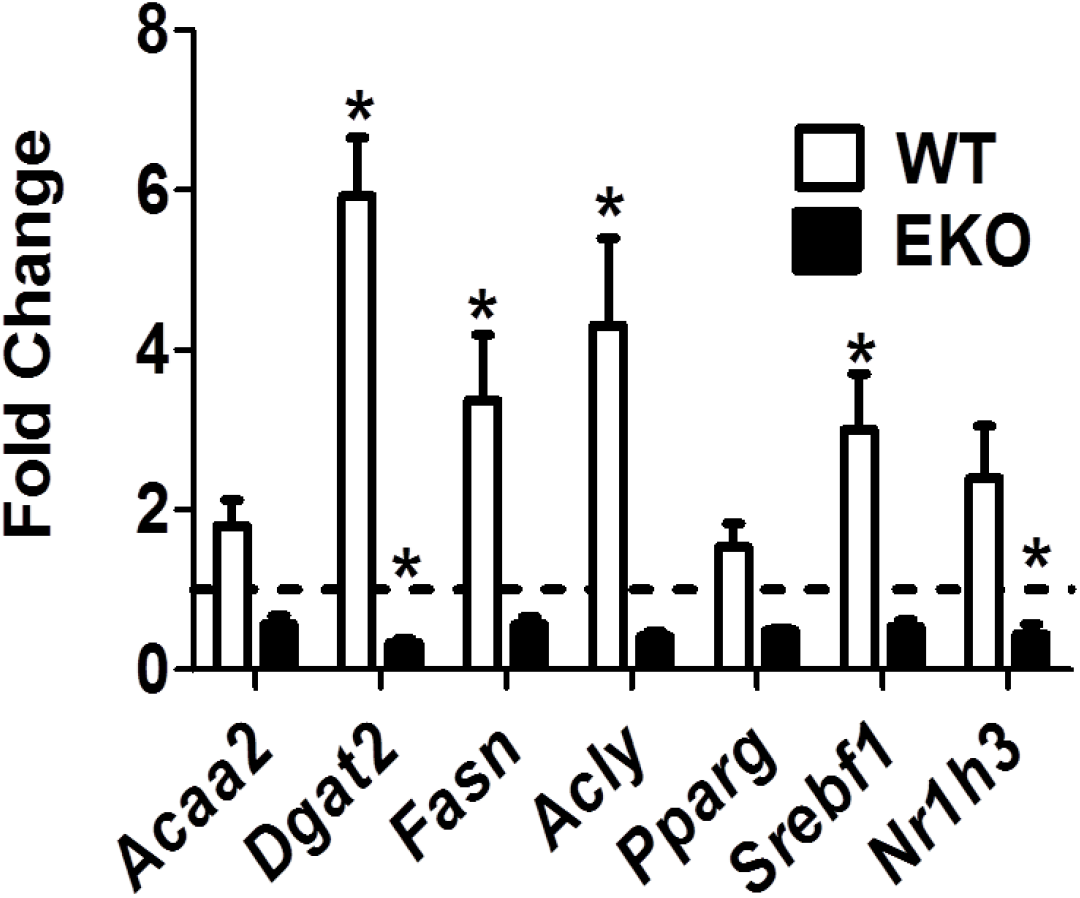
mRNA expression of genes encoding key transcription factors and metabolic enzymes which regulate fatty acid metabolism are up-regulated in WT but not EKO HFD adipose tissue. qPCR for select genes was performed on RNA isolated from epididymal VAT of normal and HFD fed WT and EKO mice. Fold expression is in comparison to ND controls of the respective genotype. Dashed line is the mean expression in ND controls. *p<0.05 vs. ND control using Student’s t-test, N=3 per group. *Acaa2*=Acetyl coA acyltransferase 2, *Dgat2*=Diacylglycerol O acyltransferase 2, *Fasn*=Fatty acid synthase, *Acly*=ATP citrate lyase, *Pparg*=Peroxisome-proliferator activated receptor gamma, *Srebf1*=Sterol response element binding factor 1, *Nr1h3*=Nuclear receptor sub family 1, group H, member 3

To gain a more comprehensive picture of these changes in gene expression, we performed whole genome expression profiling of VAT from WT and EKO mice fed ND and HFD for 24 weeks. Using the Affymetrix Mouse Gene 1.0 ST chip, we performed 4 biological replicates per condition. We defined differential expression as a fold change >1.5 at an alpha of p<0.05. When comparing ND WT to HFD WT adipose tissue, expression of 880 genes was significantly different between conditions (3% of represented probes)(Figure 5, Supp. Table 1). In contrast to WT VAT, EKO VAT was remarkably insensitive to dietary fat in terms of quantitative changes in gene expression. Expression of only 53 genes was differentially regulated in EKO adipose tissue when comparing ND versus HFD (0.1% of represented probes)(Figure 5, Supp. Table 1). Gene set enrichment analysis (GSEA) revealed that the top ten molecular pathways represented in the over expressed genes in WT HFD mice were all inflammatory signaling nodes (Figure 5). Interestingly, the pathway with the greatest fold enrichment in the EKO HFD fed mice was the peroxisome proliferator activated receptor alpha (PPARα) pathway; less than half of the remaining pathways represented in the EKO HFD mice were inflammatory pathways (Figure 5).

**Figure 5.**
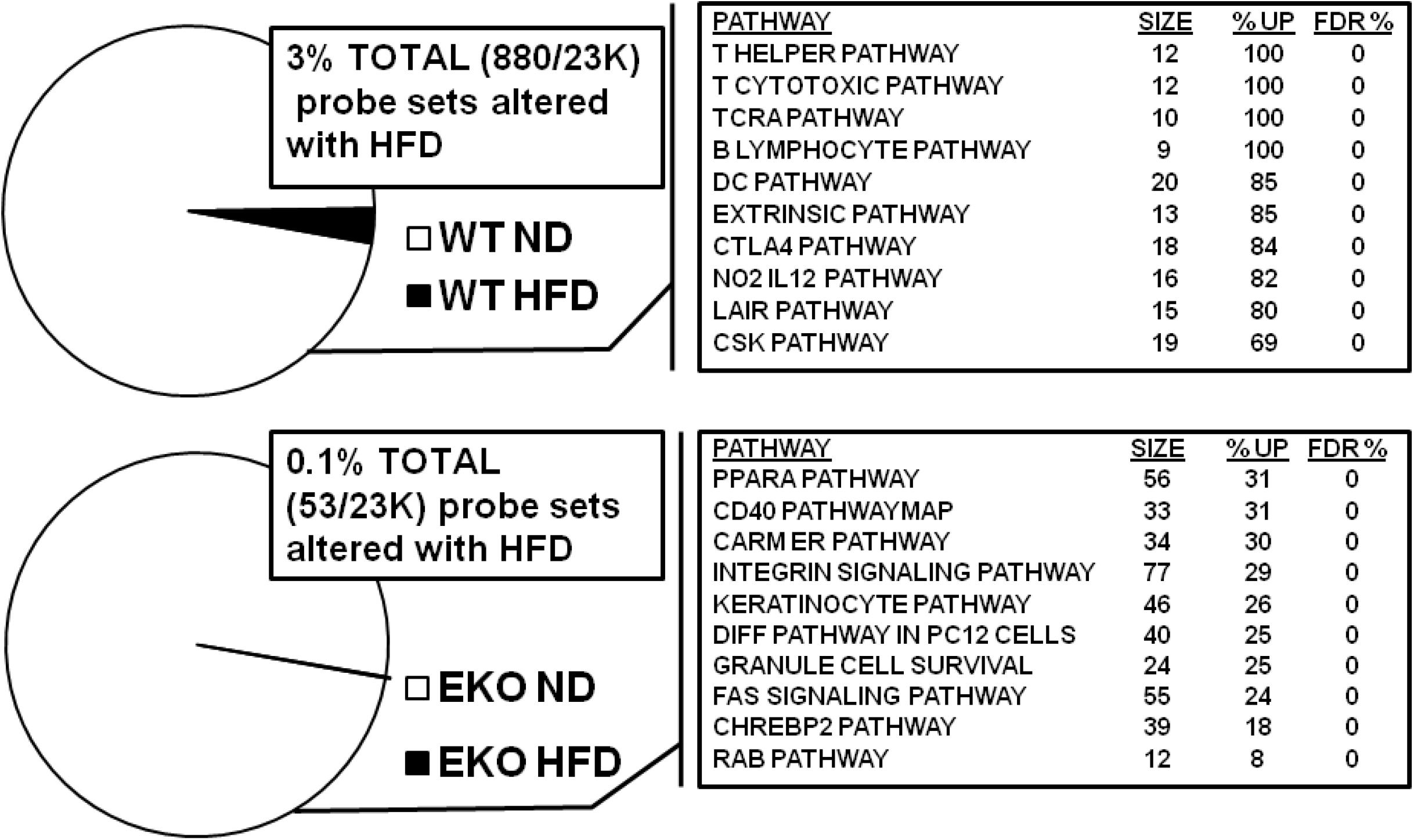
Comparative genomics reveal that gene expression in EKO VAT is relatively insensitive to HFD. 3% of represented probes were differentially expressed in WT HFD mice (top), while only 0.1% of total probes were differentially expressed in EKO HFD mice (bottom). GSEA showed that the top ten most enriched pathways in WT HFD mice were all involved in inflammatory pathways (top box). In contrast, the most highly represented pathway in EKO HFD mice was the PPARα signaling family (bottom box).

We then specifically queried expression of an *a priori* list of genes whose products encode proteins which function in fatty acid metabolism, including PPARγ target genes, pro-adipogenesis genes, β oxidation genes, and cholesterol metabolism genes (Table 3). PPARγ target genes such as insulin receptor substrate 1 (*Irs1*), Complement component d (*Cfd*), facilitated glucose transporter member 4 (*Glut4/Slc2a4*), and adiponectin (*Adipoq*) were uniformly and significantly down regulated in response to HFD in WT mice (p<0.05)(Table 3). In contrast, PPARγ target genes were not significantly down-regulated in EKO adipose tissue. Interestingly, genes belonging to cholesterol metabolism pathways were significantly up-regulated in the HFD fed WT mice, whereas there was no response to HFD in EKO adipose tissue. The only gene set consistently up-regulated in HFD fed EKO mice were those encoding enzymes responsible for fatty acid catabolism and β oxidation (Table 3). These findings corroborated the GSEA, which showed increased expression of genes in the PPARα signalling pathway (Figure 5).

**Table 3.**
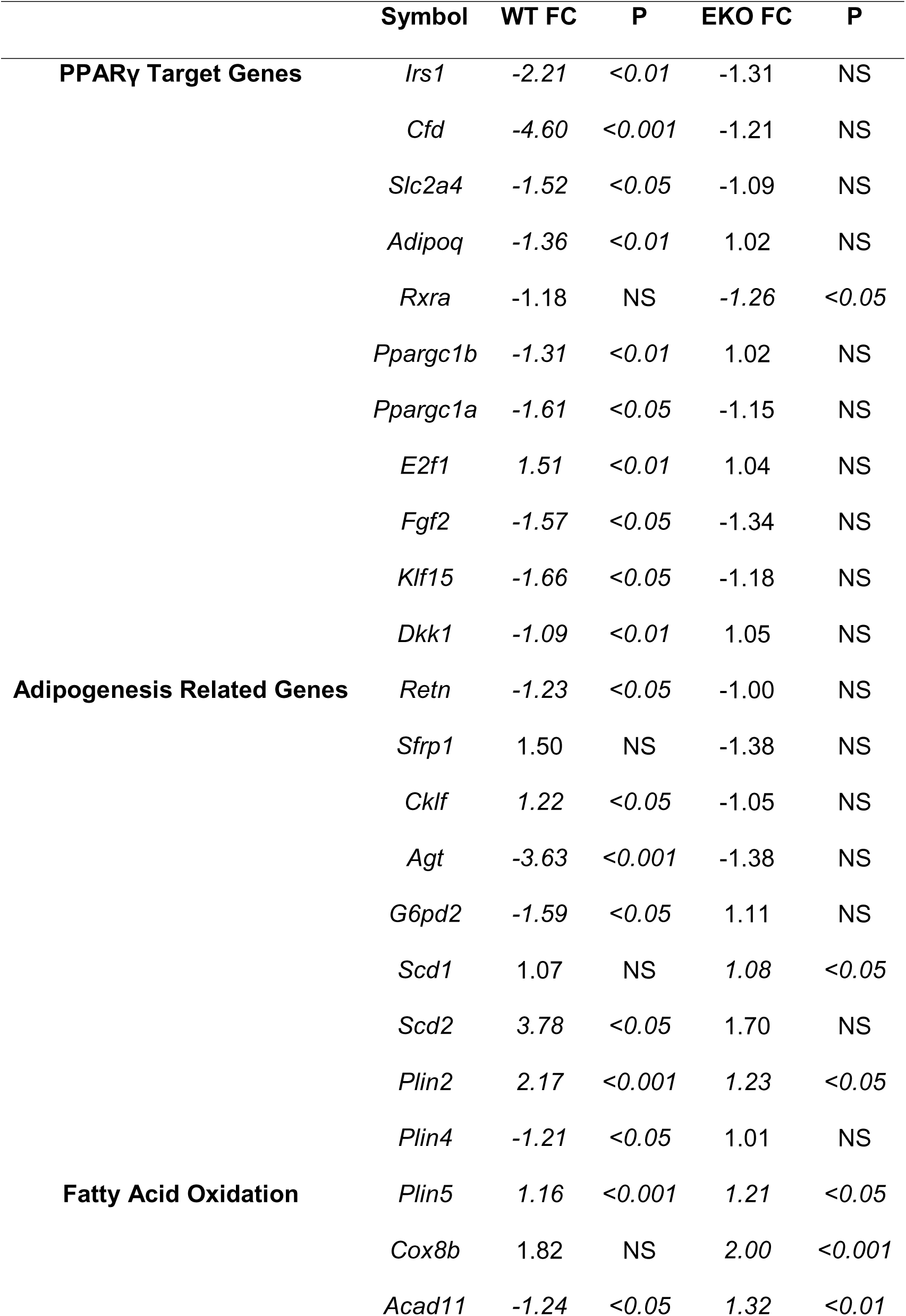
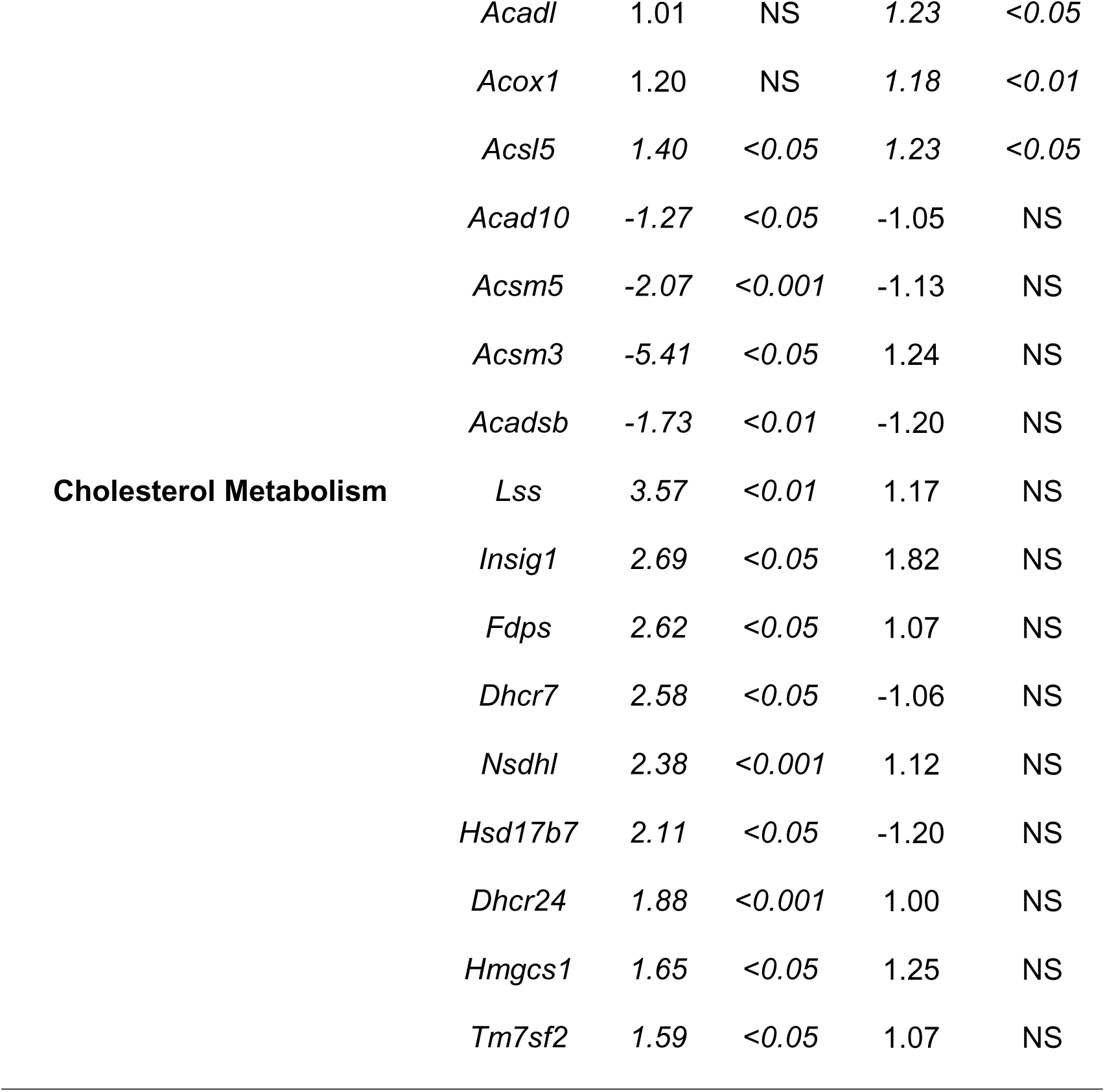
Differential Regulation of Select Metabolic Gene Sets in Response to HFD in WT vs. EKO

### Reduced F480 staining and Macrophage Specific Gene Expression EKO HFD VAT

Considering the dramatic increase in expression of inflammatory genes in the WT HFD VAT (Figure 5, Supp. Table 1) we decided to confirm the extent of macrophage infiltration with HFD by immuno-histochemistry for the macrophage marker F480+ (Figure 6). WT HFD VAT had abundant macrophage F480+ staining, with most appearing as large multi-nucleate cells in crown like structures (CLS)(Figure 6B). In contrast, there were very few F480+ macrophages or CLS in the EKO HFD VAT (Figure 6D). We then queried our microarray analysis for expression of genes known to be enriched in macrophages and immune cells (Figure 6E). WT HFD mice had a significant increase in expression of the macrophage specific genes *Emr1* (2 fold), *Cd68* (3 fold), and *Itgax* (5 fold), whereas there was no increase in EKO HFD fed mice (Figure 6E). Finally, we validated expression of immune specific genes shown to be up-regulated in the VAT of WT HFD mice on our microarrays using RT-qPCR (Figure 6F). Expression of the macrophage specific genes *Emr1, Cd68*, and *Itgax* were 8, 10, and 40 fold increased in the WT HFD VAT compared to WT ND mice (p<0.05). In contrast, there was no increase in expression in the HFD fed EKO mice. Expression of *Ccl2, IL7r*, and *Tnfα* was also significantly increased in the WT HFD VAT (p<0.05) (Figure 6F).

**Figure 6.**
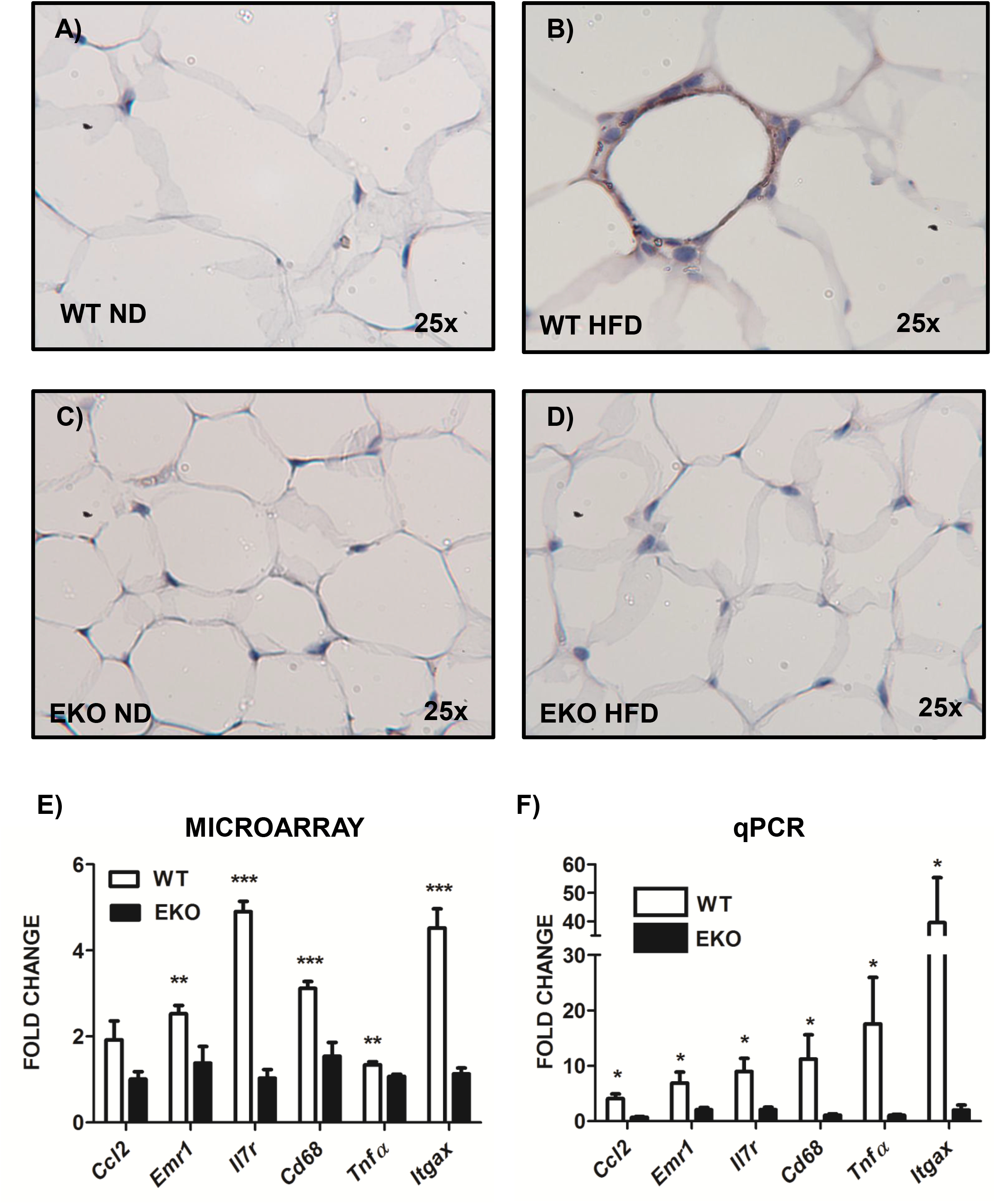
WT HFD VAT has greater F480+ Macrophage staining and expression of macrophage and immune cell specific genes. (A-D) Epididymal VAT from WT and EKO mice fed ND and HFD for 24 weeks was harvested and stained with the macrophage specific marker F480+. WT adipocytes were larger than EKO at baseline and with HFD and contained greater numbers of F480+ cells (B). (D) Microarrays revealed that expression of macrophage and immune cell specific genes were highly up regulated in WT but not EKO HFD mice. *p<0.05 vs. WT ND using the Student’s T test. (E) qPCR validation of immune cell enriched genes shown to be up regulated in the microarray analyses. (n=6 per group). *p<0.05 vs. WT ND using the Student’s T test

## Discussion

It is now well established that the EKO mouse is resistant to DIO and remains glucose tolerant when compared to WT HFD fed controls^6–8, 10, 23, 24^. Our results confirm these prior studies and make three novel contributions: 1) post-prandial VO2 (ml/hr) is increased in HFD fed EKO mice, 2) serum β-hydroxybutyrate concentrations are increased in EKO mice, and 3) the VAT of EKO mice is comparatively insensitive to HFD in terms of global alterations in gene expression.

Regarding our studies of whole body metabolism, compared to WT controls, EKO mice had a significantly reduced fat mass in response to HFD; this was apparent within 8 weeks of starting a HFD, and continued until 24 and 38 weeks (Supplemental Fig 1). WT mice had a 5 fold increase in fat mass after 24 weeks HFD while EKO mice had only a 2 fold increase (Table 1). Huang *et al*. have shown that endogenous adipocyte apoE is required for both LPL dependent and VLDR endocytic uptake of triglyceride rich lipoprotein particles, and this would account for reduced fat mass ^11^. However, how do we account for the apparent caloric imbalance in EKO mice?

We detected no differences in food intake or physical activity between WT and EKO animals (Table 1). Others have shown that intestinal absorption of lipid is normal in the EKO mouse^25^. Therefore, we hypothesized that EKO mice might have increased energy expenditure. Gao *et al*. found that EKO mice on the Ay/+ background had increased VO2 in light and dark phases ^7^. In contrast, Hofmann *et al.* found no differences in energy expenditure after 16 weeks of HFD ^8^. Indeed, we found that when VO2 was normalized to body weight (ml/hr/per kg body weight) HFD fed EKO mice had greater rates of VO2 (Table 1, Figure 2A). However, when we analyzed raw or non-normalized VO2 (ml/hr), there was no significant difference between groups (Table 1, Figure 2B). ANCOVA revealed that diet, but not genotype, was significantly associated with VO2 (ml/hr) and EE (kcal/hr). The linear model incorporated weight, lean body mass, and fat mass as covariates to account for these independent variables (Table 2). Our use of non-normalized measures of energy expenditure and ANCOVA are in conformity with a recent consensus statement by an expert panel of scientists^26^. It is likely that prior studies which included body weight in the estimation of VO2 (ml/hr/kg) and EE (kcal/hr/kg) erroneously ascribed a greater energy expenditure to EKO mice. Despite these issues, we did find that non-normalized post-prandial VO2 (ml/hr) was significantly greater in HFD EKO mice than HFD WT mice, a factor which, over the life of the animal, may contribute to reduced weight gain (Figure 2C,D)^27^.

Consistent with prior reports, we found that the livers of HFD fed EKO mice were resistant to hepatosteatosis (Figure 3)^6, 7, 24, 25^. As apoE is a ligand for the LDL, VLDL, and remnant receptor (LRP1), the liver cannot take up chylomicron remnants and LDL and therefore may be less prone to lipotoxicity. This highlights the important contribution of dietary fat to hepatic steatosis. However, as VLDL secretion is also dependent upon apoE, and rates of hepatic VLDL secretion are decreased in EKO mice, one might expect hepatic TG to accumulate despite reduced uptake ^28, 29^. We observed the exact opposite, that EKO mice were protected from hepatosteatosis with HFD. Increased hepatic β-oxidation of fatty acids may reduce liver TG accumulation in EKO mice ^25^. The increased plasma β-hydroxybutyrate concentrations that we observed in ND and HFD EKO mice are consistent with this idea (Table 1).

Perhaps the most dramatic findings of our study stem from the microarray analyses. (Figure 5, Table 3, & Supp. Tables 1) HFD feeding in EKO mice resulted in significantly altered expression of only 0.1% of probe sets; in contrast, WT HFD mice had a 30 fold greater number of genes whose expression was altered in response to HFD (3%). We initially hypothesized that EKO adipocytes may compensate for decreased TG uptake by increasing *de novo* fatty acid synthesis. qPCR showed that genes encoding lipogenic enzymes (*Acly, Fasn, Dgat*) were up-regulated in WT HFD adipocytes but not in EKO adipocytes (Fig 4). We then queried the microarray data for an *a priori* determined list of genes involved in fatty acid metabolism. We again failed to detect an increase in expression of the same lipogenic genes in EKO adipocytes. However, we did find some interesting patterns of expression which correlated with our metabolic studies (Table 3).

First, important PPARγ target genes such as *Adipoq, Cfd, Irs1 and Glut4/Slc2a4*, were all down regulated in response to HFD in WT VAT; this correlated with the potent inflammatory response in this tissue and impaired whole body glucose metabolism (Table 3). Inflammation, specifically TNFα, is thought to reduce PPARγ activity in adipocytes ^30^. Second, a panel of genes encoding proteins in the cholesterol biosynthesis pathway (*Hmgcs1, Insig1, Lss, Hsd17b7*) was up regulated in WT HFD fed VAT (Table 3). It has been shown in previous studies that large adipocytes are cholesterol deficient due to their requirement for a large plasma membrane, therefore they activate expression of genes to increase cellular cholesterol ^31^. In that regard, it is also notable that the expression of the VLDL (1.4 fold, p<0.05) and LDL (1.8 fold, p=NS) receptors was increased (1.4 fold) in the WT HFD mice, likely as an additional mechanism to increase cellular cholesterol (Supp Table 1). Cholesterol deficiency causes cellular insulin resistance by reducing signaling through caveolin 2 enriched lipid rafts ^31^. Of note, the diameter of WT type adipocytes was much greater than EKO adipocytes at baseline and especially with HFD (Fig 6).

In regards to the increased numbers of macrophages in the WT HFD VAT, the microarray data indicate that multiple mechanisms of macrophage lipid uptake appear to be activated, all of which parallel foam cell formation in the atherosclerotic plaque. For example, enzymatically modified LDL particles are internalized mainly by type 1 (opsonin) and type 2 (complement) mediated phagocytosis ^32^. Expression of both *Cd16/Fcgr3*, which binds C reactive protein and IgG coated LDL particles, and *Cd11b*, which binds complement coated LDL particles, is increased in WT HFD VAT (1.48 and 1.71 fold respectively p<0.05, Suppl. Table 1). Type 1 and type 2 phagocytosis lead to the formation of large, membrane associated lipid droplets ^32^. In contrast, expression of the macrophage scavenger receptor *Msr1*, which is responsible for receptor mediated of predominately oxidized LDL particles, is also increased (1.92 fold, p<0.05, Suppl. Table 1). Expression of these genes is not altered in EKO VAT with HFD. We do not suggest that modified and oxidized LDL are necessarily the ligands for macrophages in adipose tissue, only that it is likely that there are multiple complex lipid antigens released from dying (or viable) adipocytes in WT HFD VAT which broadly stimulate the innate immune response. The decreased cell size of EKO HFD adipocytes may prevent cellular stress and death thus preventing this inflammatory response.

That we do not see increased macrophages in the VAT of HFD EKO mice is not inconsistent with the results of others who have shown that 1) primary EKO adipocytes have increased rates of lipolysis *ex vivo* and 2) FFA release is a stimulus for macrophage infiltration ^5, 17^. On a cellular basis, because the lipolytic rate depends on cell surface area (4πr2), and EKO adipocytes are almost ½ the diameter of WT adipocytes, FFA release per cell is actually likely much smaller in EKO adipocytes. Second, rates of lipid oxidation may be increased in EKO adipocytes, resulting in less release of FFA from hydrolyzed TG. Finally, macrophage infiltration in response to lipolysis has been shown in obese animals subjected to fasting, thus, in negative energy balance, which was not the case for EKO mice ^17^.

In regard to changes in gene expression in EKO adipose tissue, expression of genes encoding enzymes important for fatty acid oxidation were broadly up-regulated, for example, *Oxpat/Plin5, Cox8b, Acsl5, Acox1*, and *Acadl* (Table 3). Many of the same genes were down regulated in WT HFD VAT. Interestingly, the gene with the highest fold increase in expression in EKO HFD VAT, lactase-like (*Lctl*), is a recently described member of the fibroblast growth factor family which is specifically expressed only in brown adipose tissue ^33^(Supplemental Table 1). Overall, these findings correlated with the results of GSEA, in which the PPARα signaling family was top represented pathway in EKO HFD fed mice (Figure 5). Whether or not fatty acid oxidation is actually functionally increased in EKO adipocytes, and relates to their decreased cell size, will require further study.

In summary, using a combination of physiologic studies, genomics, and immunohisto-chemistry, we describe novel features of adipocyte metabolism in EKO mice which will provide a starting point for future studies. We confirmed the observation that, despite severe hyperlipidemia, EKO mice fed a HFD are leaner and insulin sensitive than their WT littermates. This may be due to failure of adipose, skeletal muscle, and liver to properly metabolize circulating triglyceride rich lipoproteins. Second, using careful statistical analysis of metabolic cage data in accordance with expert guidelines, we show that post-prandial VO2 (ml/hr), but not total VO2 or EE is increased in EKO mice. This may partially account for their lean phenotype. Finally, using the first whole genome comparison of gene expression of WT and EKO adipose tissue, we show that EKO mice have decreased expression of inflammatory genes, and increased expression of genes encoding enzymes responsible for lipid oxidation. These results suggest that limiting adipocyte size or TG uptake may be a viable therapeutic strategy, provided that peripheral mechanisms to prevent atherogenic hyperlipidemia can be activated.

## Supporting information

## Acknowedgements

We thank the Morphology Core Facility and the Genomics Core Facility of the UMASS Medical School Diabetes and Endocrinology Research Center (DK-32520) for assistance with histology and microarray analyses, respectively. These studies were supported by National Institute of Diabetes and Digestive and Kidney Diseases Grants DK-30898 and DK-60837 (to M. P. Czech), American Heart Association Predoctoral Fellowship Grant No. 11PRE5000000 (to T. P. Fitzgibbons), and the National Mouse Metabolic Phenotyping Center at UMass sponsored by the National Institutes of Health, National Institute of Diabetes and Digestive and Kidney Diseases, and National Heart, Lung, and Blood Institute NIH Grant 5U2C-DK093000 (to J.K. Kim).

---

**SUPPLEMENTARY FIGURE 1.**
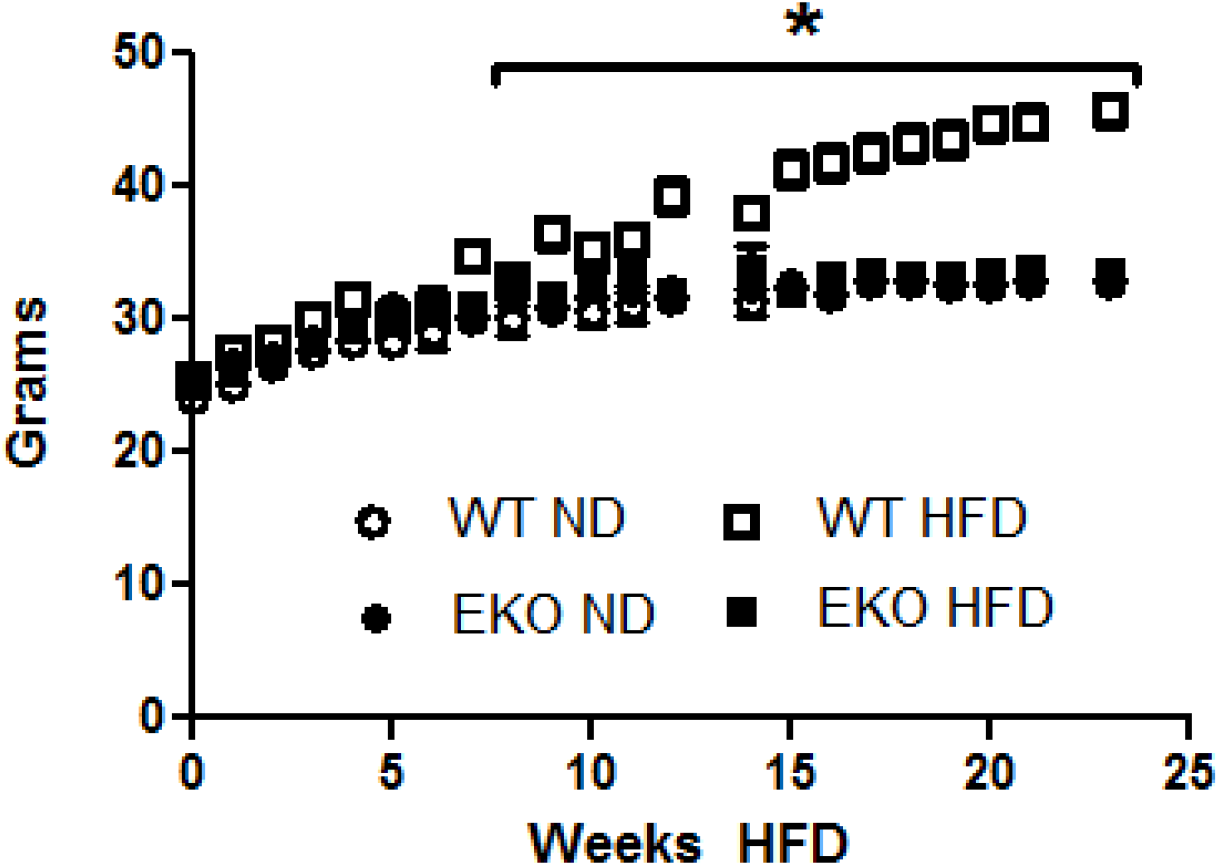

**SUPPLEMENTARY FIGURE 2.**
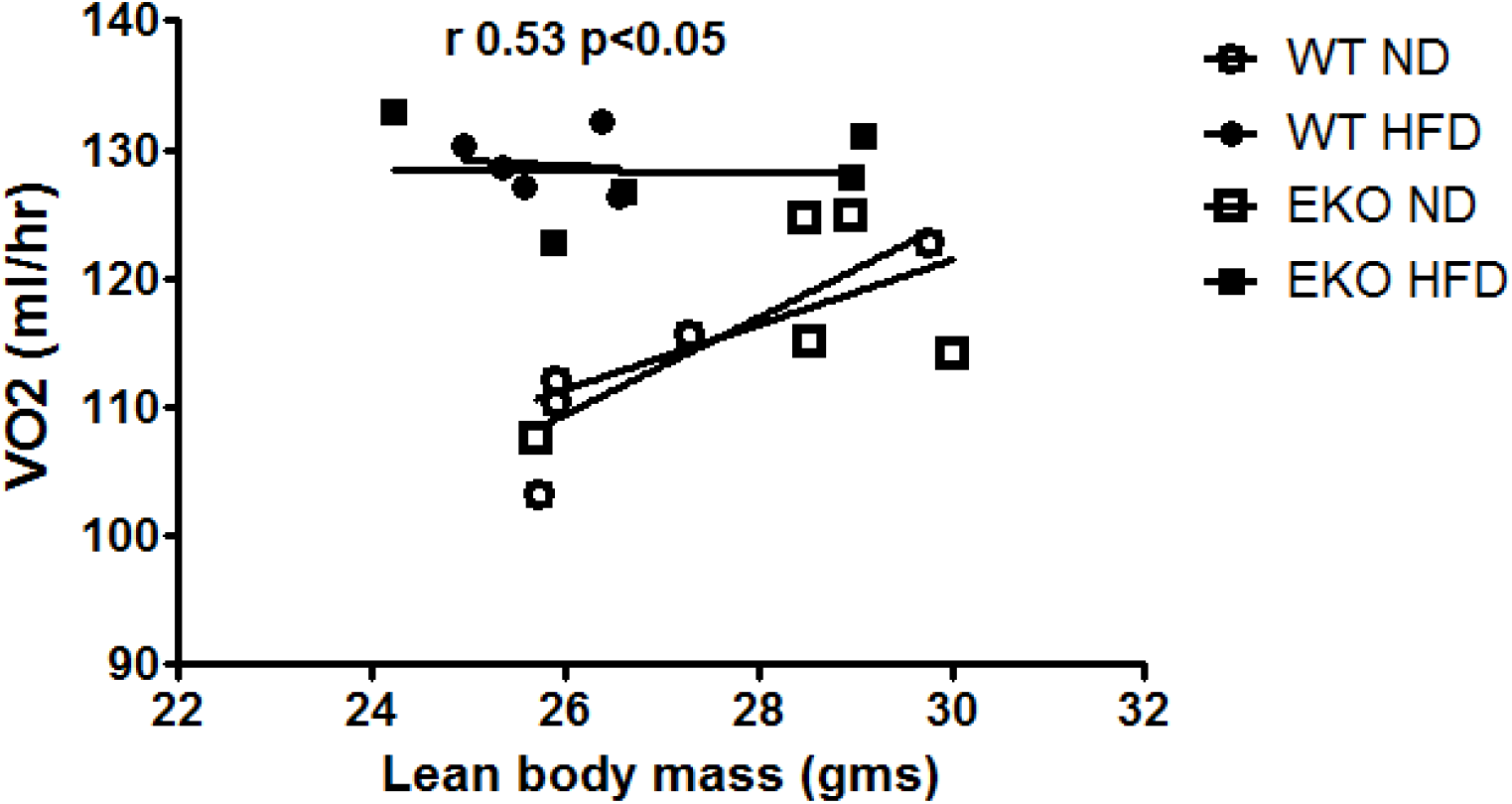

